# Quantifying the binding affinity of a pharmacological chaperone to transient unfolded states of a normally folded protein

**DOI:** 10.64898/2026.08.27.747683

**Authors:** Shubhadeep Patra, Craig R. Garen, Michael T. Woodside

## Abstract

Binding of ligands to partially or fully unfolded proteins can play a key role in the mechanism of cellular and pharmacological chaperones, facilitating proper folding. However, it is challenging to quantify the binding affinity of ligands for unfolded states in a protein that is normally folded, as the methods standardly used to destabilize the native fold also affect ligand binding. We used single-molecule force spectroscopy to unfold single protein molecules without altering solution conditions and observe interactions of a ligand with unfolded states. Focusing on pentosan polysulfate (PPS), an anti-prion pharmacological chaperone previously shown to interact with both the native and partially or fully unfolded states of the prion protein, we measured the concentration-dependent effects of PPS binding on the conformational dynamics of bank vole prion protein (BvPrP) molecules held in optical tweezers. We found that PPS stabilized certain partially unfolded intermediate states of BvPrP as well as the fully unfolded state. Strikingly, the tendency to bind unfolded states instead of the folded state increased as the PPS concentration was reduced, implying a higher affinity to unfolded states. From the relative amount of binding to unfolded versus folded states, we estimated that PPS bound roughly 100-fold more tightly to unfolded states than to the native state of PrP. These results reinforce the likely importance of unfolded states in prion misfolding and propagation. More generally, they show how binding affinity to transient, unstable states can be estimated.

## INTRODUCTION

Natively structured proteins generally tend to remain in their folded state, as it is the conformation that supports their native function. However, many proteins are unable to fold spontaneously, requiring assistance when still fully or partially unfolded to find the correct structure and avoid misfolding. Furthermore, various conditions including mutations, environmental stress, or cellular crowding can induce folded proteins to undergo partial or complete unfolding, leading both to the loss of function and to the possible triggering of misfolding and nonspecific aggregation^1–4^. In the cell, molecular chaperones help avoid these outcomes, interacting preferentially with unfolded or partially folded proteins to assist in folding, prevent aggregation, and/or restore functional activity^5–8^. Pharmacological chaperones that mimic the action of molecular chaperones to help prevent misfolding have also been developed, as a therapeutic strategy for diseases including cystic fibrosis, lysosomal storage disorders, cancer, and neurodegenerative disorders^9–11^, with many small-molecule chaperones binding to partially unfolded or misfolded intermediates^12–15^. Studying the interactions of ligands with partially or fully unfolded conformations of proteins that are natively folded is therefore critical for understanding both the fundamental science of protein misfolding/aggregation, identifying potential therapeutic targets—including alternative druggable sites that are hidden in the native fold—and probing the mechanism of action of chaperones.

Interactions with ligands and unfolded states have been studied extensively in proteins that are intrinsically disordered. Because such proteins populate unfolded states under native conditions, a wide range of methods can be used to characterize ligand binding kinetics, thermodynamics, stoichiometry, and structure, including NMR, calorimetry, surface plasmon resonance, and ensemble or single-molecule fluorescence spectroscopies^16–18^. However, it is more challenging to do the same for interactions with unfolded states in proteins that are normally folded, because the unfolded states are typically unstable and transient, and they often represent off-pathway or stress-induced states of the natively folded protein rather than functionally regulated conformations as in intrinsically disordered proteins. In the absence of conditions destabilizing the native fold, unfolded states are sufficiently rare that they are likely to escape detection by conventional binding assays. However, the methods generally used to destabilize the folded state (*e*.*g*., temperature or pressure changes, chemical denaturants, pH or ionic strength changes,…) affect the protein globally and hence also alter the interactions that stabilize ligand binding, making quantification of the interaction between the ligand and unfolded protein difficult to measure and interpret.

One approach that overcomes these challenges is single-molecule force spectroscopy (SMFS), wherein force is applied individually to a particular protein molecule in order to destabilize its structure^19^. Because the force acts locally, leaving all other molecules in solution and the global solution conditions unaffected, the unfolded states generated by the applied force are able to interact with ligands more similarly to how they would normally do so. By analyzing how the signatures for unfolding/refolding that characterize the conformational dynamics of the protein, including changes in the thermodynamic stability, force distributions, kinetics, intermediates, and pathways^19,20^, change in the presence of the ligand, binding to fully or partially unfolded states can be detected and distinguished from binding to the native state^21–24^.

To demonstrate quantification of ligand binding to unfolded states using SMFS, we focused on anti-prion pharmacological chaperones that bind to the mammalian prion protein, PrP. Misfolded PrP can propagate by converting natively folded molecules to the mis-folded isoform, generating fatal, transmissible neurodegenerative diseases in a wide range of mammals, including scrapie in sheep, chronic wasting disease in cervids, and Creutzfeldt–Jakob disease in humans^25,26^. Similar prion-like propagation of misfolding has since been observed in several other proteins linked to a wide range of neurodegenerative disorders in humans, including Alzheimer’s and Parkinson’s diseases, amyotrophic lateral sclerosis, and frontotemporal dementia^27^. Efforts to find therapeutics that inhibit misfolding and conversion^28–33^ have discovered a number of small-molecule pharmacological chaperones^34–36^. In addition to interacting with natively folded PrP^34,37,38^, some of these compounds have been reported to interact with partially or fully unfolded states of PrP^39,40^, similar to the action of molecular chaperones^5,21,41^. Such interactions could be crucial, as the transmissible misfolded isoform of PrP contains no remnants of native structure and therefore requires complete reconfiguration of the native fold via partially or fully unfolded states. However, the interactions of such anti-prion compounds with unfolded states of PrP remain poorly characterized: for example, the binding affinity to partially and fully unfolded states is unknown, and it is unclear if the ligands bind to partially native or non-native intermediates. Understanding the interactions of pharmacological chaperones with unfolded PrP in more detail should therefore help improve our understanding of anti-prion action.

Here, we have investigated the binding of one of such anti-prion chaperone, pentosan polysulfate (PPS) (Fig. 1), to bank vole PrP (BvPrP). Bank voles are “universal receptors” for prion diseases, having no barrier to infection by prions from other species^42^; BvPrP folds/unfolds via multiple intermediate states and forms long-lived metastable misfolded states even in monomers, unlike other mammalian PrPs^43–45^. Measuring BvPrP folding and unfolding in optical tweezers at different concentrations of PPS, we found that PPS binds and stabilizes partially folded intermediates along the native folding pathway, as well as the fully unfolded state, and also stabilized additional intermediates not seen in the absence of PPS. By comparing the relative amounts of PPS binding to folded and unfolded states, we also quantified the binding affinity to unfolded states, finding that it is about 100 times higher than to the native fold. These results showcase a new approach for measuring the binding affinity to unfolded states, and underline the likely importance of interactions with unfolded PrP for preventing propagated misfolding in prions.

**Fig. 1.**
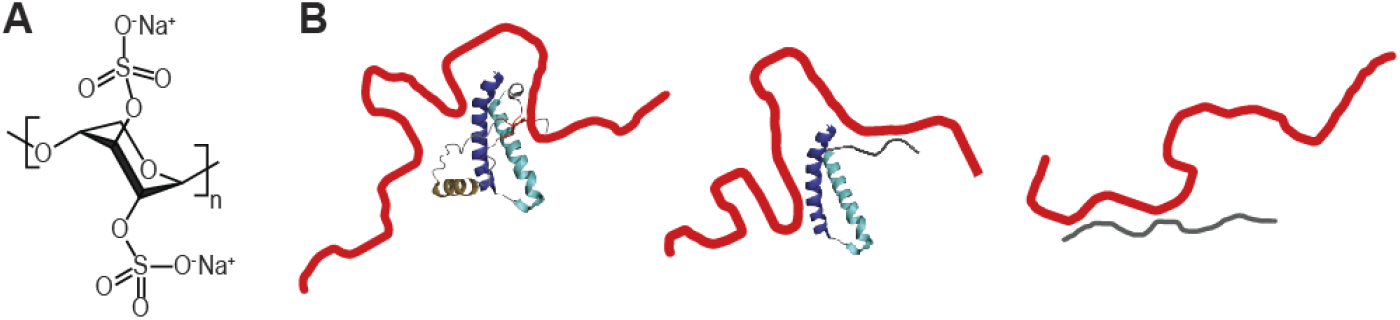
Pentosan polysulfate (PPS) as a pharmacological chaperone for PrP. (A) Structure of PPS. (B) Cartoon of PPS (red) interacting with PrP (blue) when it is natively folded (left), partially unfolded (center), and fully unfolded (right)

## RESULTS

To assess the effects of PPS on BvPrP folding, single BvPrP molecules were covalently attached to kilobase-long double-stranded DNA handles bound to polystyrene beads held in optical traps (Fig. 2A). The traps were moved apart at a constant speed to apply force on the protein and induce unfolding, then brought back together to decrease force and induce refolding. Multiple cycles of unfolding-refolding were repeated, with a wait time of 5 s at near-zero force between cycles to facilitate refolding to the native structure. Without PPS present, as reported previously, FECs typically displayed two or sometimes three discrete unfolding transitions (Fig. 2B) between ∼5–20 pN, where the extension increased abruptly concomitant with a drop in the force as segments of the structure unfolded, reflecting the formation of typically one or sometimes two intermediate states. Fitting each branch of the FEC to an extensible wormlike chain model (WLC) relating the applied force, *F*, and molecular extension, *x* (Eq. 2 in Methods), we determined the contour length (*L*_c_) of the polypeptide chain unfolded in each transition. The total change in *L*_c_summed across all transitions in these curves, Δ*L*_c_= 34.5 ±0.3 nm, matched the value expected for complete unfolding of the structured domain of BvPrP based on the NMR structure^46^. For the intermediate states, Δ*L*_c_varied between four values (Fig. 2C), reflecting four distinct intermediates (denoted I_1_through I_4_). However, roughly 1/4 of these FECs showed less than full-length unfolding, indicating that the protein did not refold properly, but rather was trapped in a misfolded state (roughly 2/3 of the cases) or stalled in one of the native intermediates (the remaining ∼1/3 of cases)^43^.

**Fig. 2.**
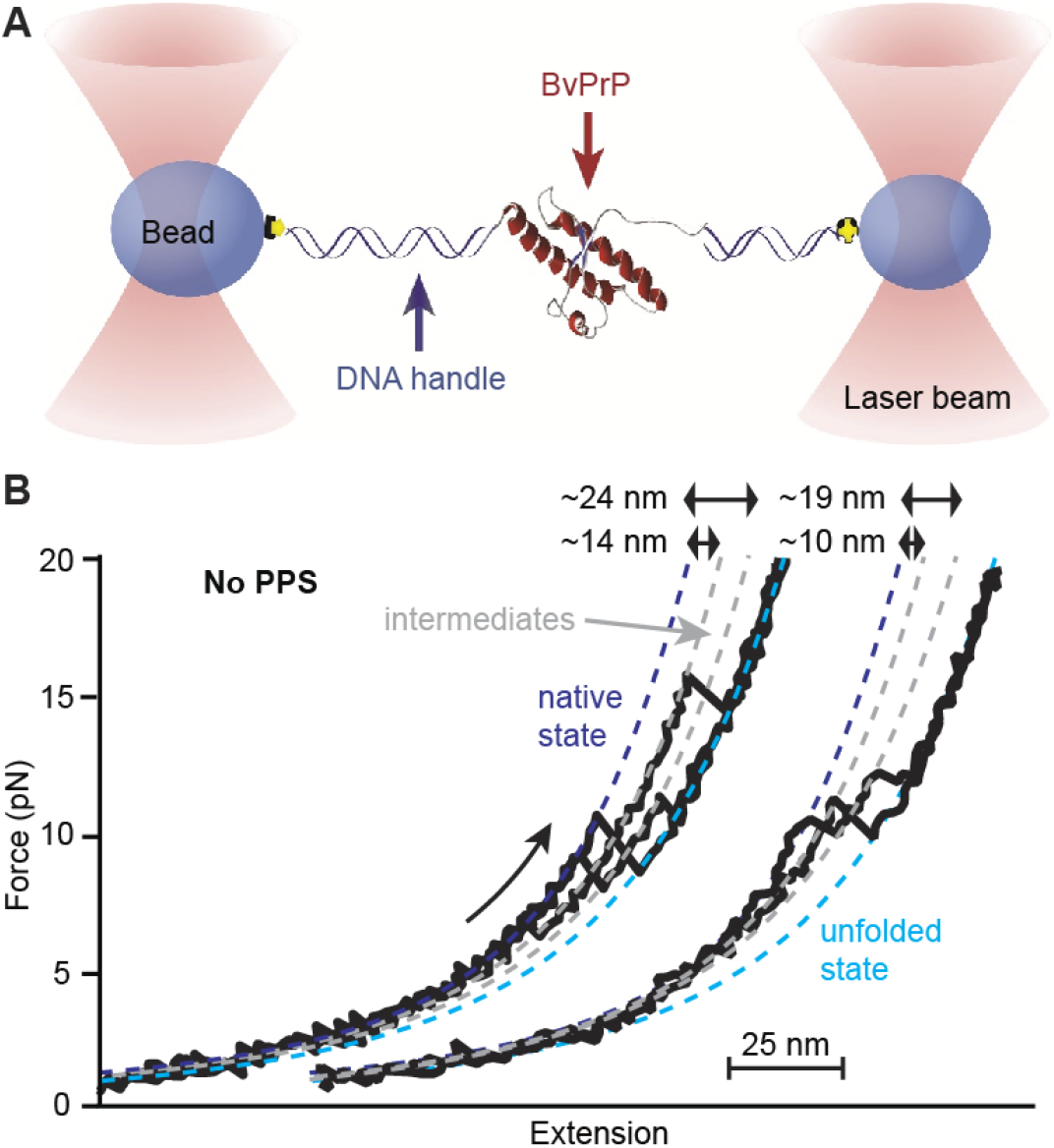
FECs of BvPrP unfolding without PPS. (A) Schematic of single-molecule force spectroscopy experiment assay. (B) FECs without PPS present show BvPrP unfolding via one or more of four distinct intermediates, with a total contour length change of 34.5 nm. Dashed lines: WLC fits to the folded (blue), intermediate (grey) and unfolded (cyan) states.

We repeated the SMFS measurements adding PPS at three different concentrations: 0.01 μM, 0.1 μM, and 0.5 μM. This range of concentrations was selected based on the affinity of PPS for the strongest binding site on native PrP, found previously by isothermal titration calorimetry (ITC) as *K*_D_= 0.19 ± 0.04 μM^39^, such that the lowest concentration was 20-fold below *K*_D_and the highest 2.5-fold above it. Adding PPS changed the FECs very noticeably (Fig. 3A). Although FECs showing complete unfolding of the native state were still observed (Fig. 3A, black), they were no longer in the majority: FECs showing length changes shorter than 34.5 nm (complete unfolding of native fold) became much more common (Fig. 3A, orange), and many curves showed no unfolding rips at all (Fig. 3A, red ). Surprisingly, the relative frequencies of FECs showing full, partial, and no unfolding varied non-monotonically with PPS concentration (Fig. 3B), with full unfolding minimal at 0.01 μM and no unfolding maximal at the same concentration, even though full unfolding was maximal and no unfolding minimal at 0 μM. These changes are somewhat counterintuitive, given that lower PPS concentration would normally be expected to lead to less-frequent binding of PPS to PrP and hence an increase in the native full-length folding, suggesting that something interesting is going on: non-native conformers of PrP must be outcompeting the native state for PPS binding.

**Fig. 3.**
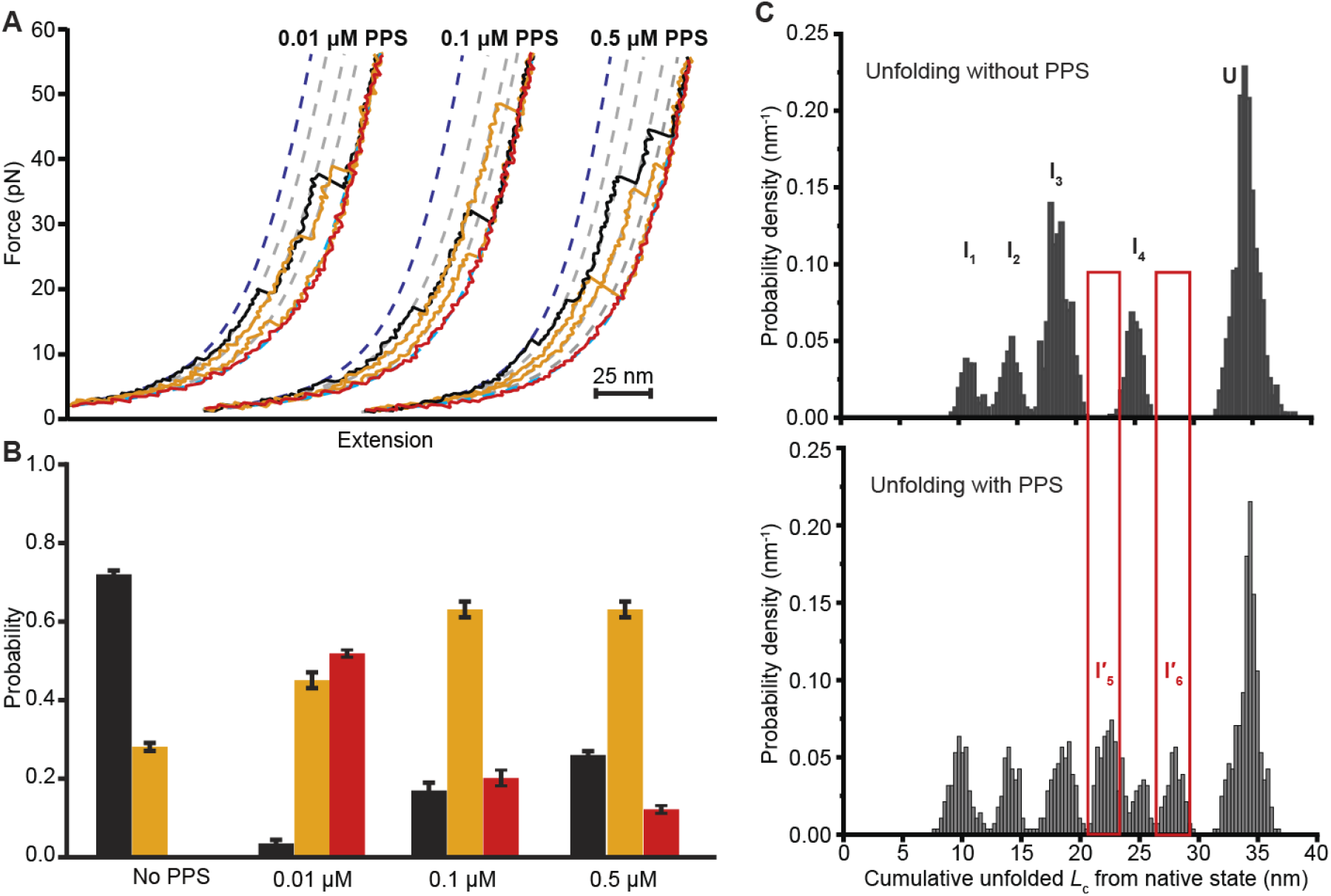
FECs of BvPrP unfolding in the presence of varying amounts of PPS. (A) BvPrP unfolds heterogeneously with PPS present at different concentrations (0.01 μM, 0.1 μM, and 0.5 μM) showing a variety of different contour length changes and wide range of unfolding forces. Some FECs show complete unfolding from native state to unfolded (black), but most show partial unfolding (orange) or no unfolding (red). (B) Fraction of curves showing full native unfolding (black), partial unfolding (orange), or no unfolding (red) as a function of PPS concentration. (C) Distribution of cumulative unfolded protein contour lengths for all states in FECs showing complete unfolding from the native state reveals that PPS binding induces the formation of 2 new intermediates and alters the populations of the intermediates present without PPS. Results from all PPS concentrations are combined together.

A second difference observed in the presence of PPS involved the intermediate states seen in the FECs. To assess these differences, we analyzed the distribution of unfolded protein contour lengths for all FECs that involved complete unfolding from the native state, combining the results from all non-zero PPS concentrations to improve the statistics. Such distributions reveal the intermediate states with distinct lengths that are occupied by the protein during the unfolding reaction and their relative abundance^47^. Comparing the distributions in the presence and absence of PPS (Fig. 3C) reveals that intermediate states were observed in the presence of PPS at the same four lengths (Table S1) as seen in the absence of PPS (I_1_at ∼11 nm, I_2_at ∼14 nm, I_3_at ∼ 18 nm, and I_4_at ∼25 nm), but with the addition of two new intermediates that were not seen in the absence of PPS: I′_5_at ∼22 nm, and I′_6_at ∼28 nm. These observations imply that PPS is not only able to bind the partially unfolded intermediates found on the native folding pathway but also stabilizes additional intermediates that are normally unstable in BvPrP. Similar behavior was previously seen for hamster PrP, where PPS binding stabilized intermediates that were otherwise normally unstable^39^.

These changes to the intermediate states imply changes to the unfolding pathways. We therefore constructed transition maps displaying all the discrete pair-wise transitions between different states observed in all FECs featuring complete unfolding from the native state, which together map out the pathways followed during unfolding. Comparing the transition maps for the unfolding of BvPrP in the absence and presence of PPS (Fig. 4), where the arrow thicknesses are proportional to the likelihood of a given transition, suggests that some of the time when PPS is present, BvPrP unfolds through the same sequence of states as seen in the absence of PPS (along the spectrum from I_1_to I_4_). However, most of the time, the unfolding is redirected to one of the two new intermediates induced by PPS binding, I′_5_and I′_6_, either entering them directly from the native state or being redirected to them from intermediates I_1_to I_3_. Regardless of which pathway is taken, PPS redirects the unfolding away from the intermediate state I_3_that dominates the unfolding in the absence of PPS.

**Fig. 4.**
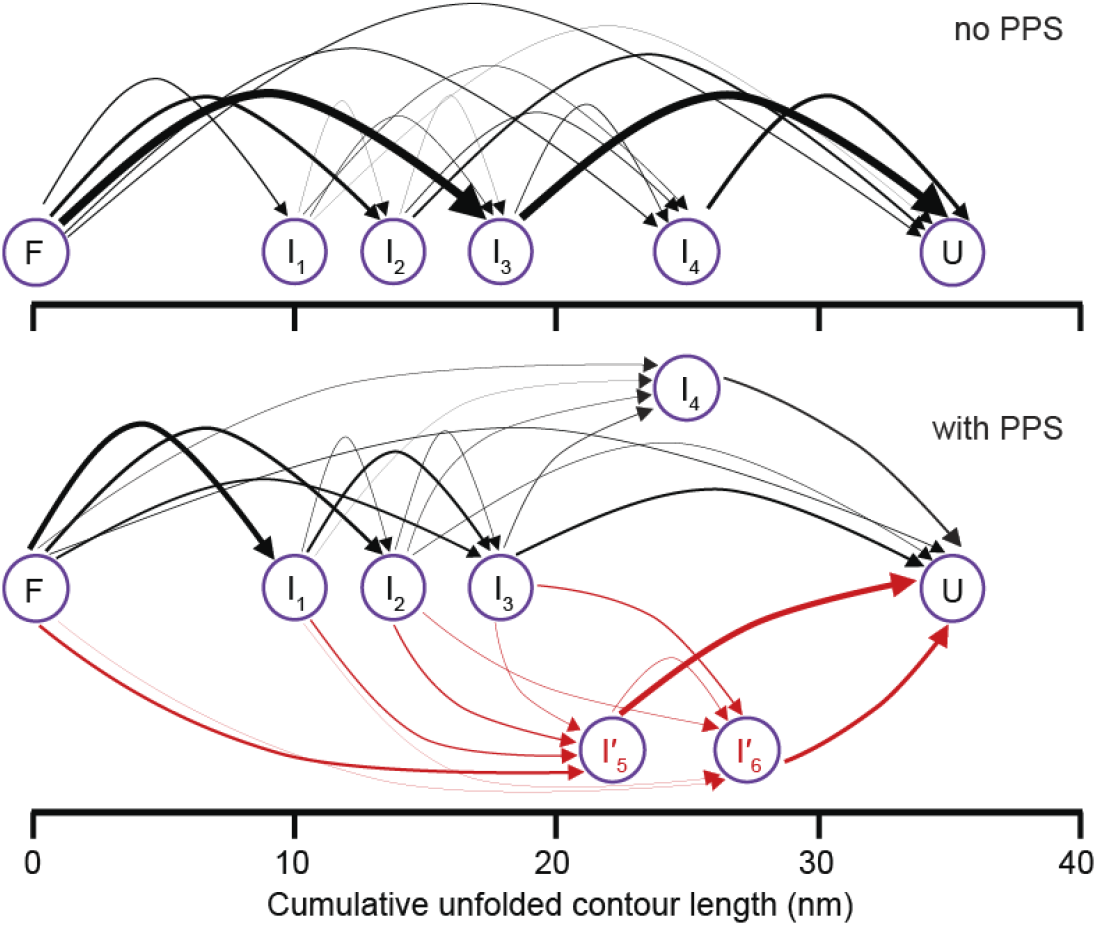
Transition maps reveal effects of PPS on unfolding pathways. Transition maps of BvPrP unfolding in the absence (top) and presence (bottom) of PPS. In the presence of PPS, only a minority of the unfolding follows pathways similar to those seen in the absence of PPS (black); most of the unfolding (red) is redirected away from these pathways to path through the new intermediates induced by PPS. Arrow thickness is proportional to transition probabilities. Only FECs showing complete unfolding from N are included in the analysis.

Yet another difference in the FECs induced by PPS binding was a noticeable increase in the unfolding forces, with the maximum unfolding force changing from ∼40 pN to over 55 pN. Examining unfolding force of the first rip, whose distribution can be modeled quantitatively in a way that subsequent unfolding events cannot^48^, we fit the PPS-dependent distributions of the first unfolding forces (Fig. 5A) to a model that accounts for heterogeneous initial states (Eq. 3)^49^, which was found previously to describe BvPrP unfolding well in the absence of PPS^43^. Both the average force (*F*_u_) and the heterogeneity (*Δ*) were highest for 0.5 μM PPS, with ⟨*F*_u_⟩ = 15.9 ± 0.3 pN and *Δ* = 12 ± 1, decreasing monotonically with PPS concentration to ⟨*F*_u_⟩ = 7.4 ± 0.2 pN and *Δ* = 3.7 ± 0.2 at 0.01 μM PPS, and in every case higher than in the absence of PPS (⟨*F*_u_⟩ = 7.0 ± 0.1 pN, *Δ* = 2.51 ± 0.08). The increased heterogeneity at higher PPS concentrations suggests that PPS binds to additional states as the concentration increases, presumably accessing binding sites with lower binding affinity.

**Fig. 5.**
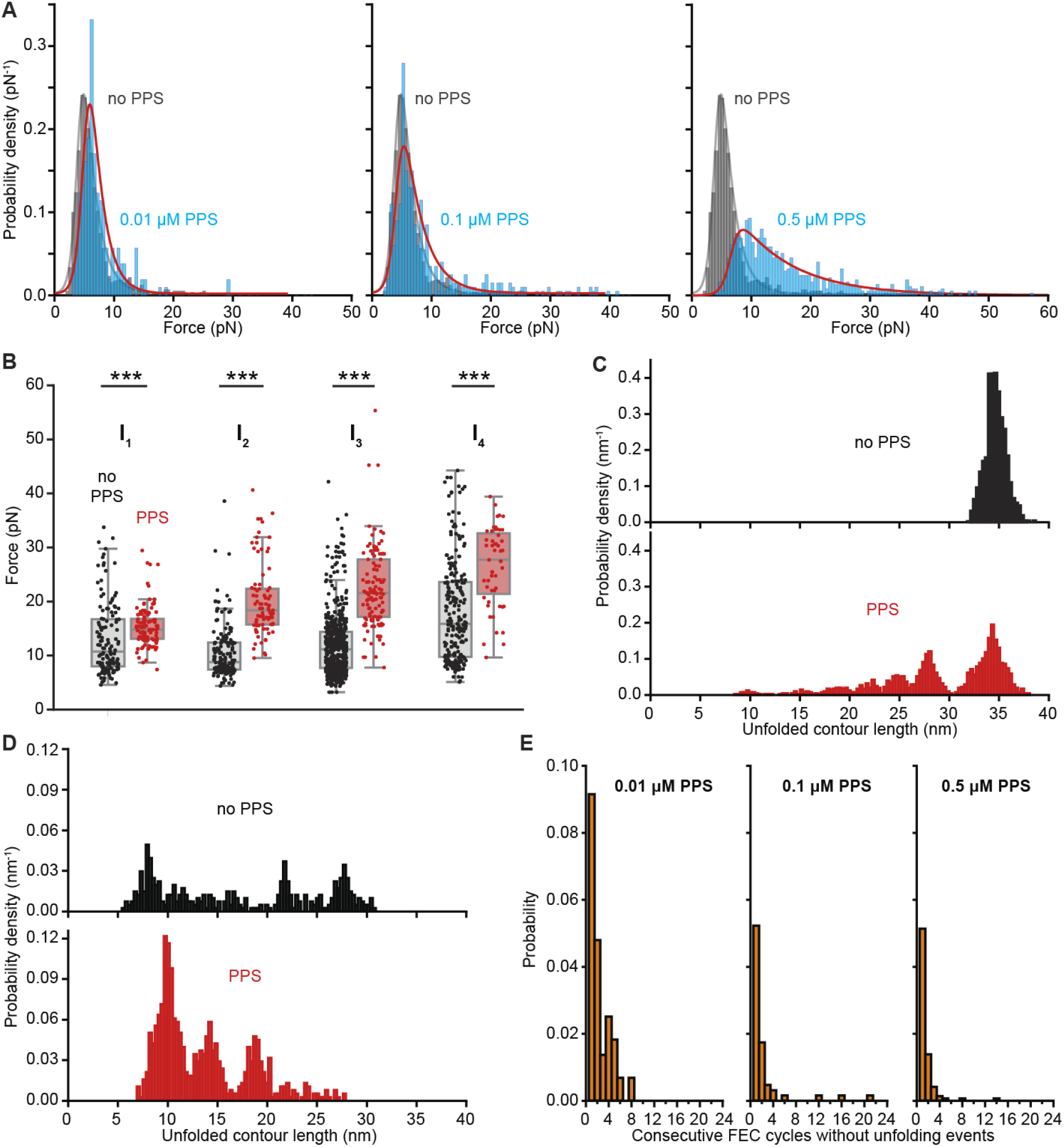
PPS binding stabilizes partially unfolded states. (A) Unfolding force distributions for the first rips in FECs show broader force ranges and higher forces in the presence of PPS (cyan) than in its absence (black). Red: fit to the heterogeneous unfolding theory (Eq. 3). (B) Unfolding forces for intermediate states I_1_to I_4_are noticeably higher with PPS present (red) than absent (black). (C) Contour length of unfolded protein at the end of each FEC, with (bottom) and without (top) PPS. (D) Contour length of unfolded protein at the beginning of each FEC, with (bottom) and without (top) PPS. (E) Probability of observing consecutive FEC cycles of varying length without any unfolding/refolding events.

Turning to the unfolding forces of the intermediate states, we focused on those that were observed in both the absence and presence of PPS (*i*.*e*., I_1_to I_4_), allowing comparisons to deduce the effects of interactions with PPS. We found that their average unfolding forces were in every case noticeably higher in the presence of PPS (Fig. 5B). PPS must therefore interact with each of these intermediates to stabilize them. Further evidence that PPS was binding to these states and stabilizing them was found from analyzing which state PrP was in at the beginning and end of each FEC, as deduced from the pattern of Δ*L*_c_values seen in curve. In the absence of PPS, every FEC ended in the fully unfolded state (Fig. 5C, top); in contrast, in the presence of PPS, FECs very frequently never reached the fully unfolded state even at the highest force applied, ending instead in a partially unfolded intermediate (Fig. 5C, bottom), implying that these intermediates were much more difficult to unfold with PPS bound. Given that these PPS-bound intermediates often did not unfold even at the maximum force reached in the FECs, the results in Fig. 5B are clearly a significant underestimate. Cataloguing the states in which FECs started (Fig. 5D), PPS again induced large changes, greatly increasing the likelihood of starting in a partially unfolded state, implying that PPS interactions with partially or fully unfolded PrP reduced the folding rate. In some cases, BvPrP only refolded partially; in others, it remained in the state in which the previous unfolding FEC ended (Fig. S1).

Indeed, in some cases the protein remained locked in the same state, without unfolding or refolding, for multiple cycles (Fig. 5E). This effect became more prominent as the PPS concentration was reduced, as with the general frequency of no-event FECs.

Lastly, we used the unfolding FECs measured at different PPS concentrations to estimate the binding affinity of PPS for partially and fully unfolded states. The binding affinity to the native fold of PrP was measured previously by ITC, revealing two independent binding sites with *K*_D1_= 0.19 ± 0.04 μM and *K*_D2_= 8 ± 2 μM^39^, implying that at the lowest concentrations used here (0.01 μM), interactions with the native state are unlikely. Nevertheless, significant effects were seen at this concentration, including a large increase in events where BvPrP did not unfold or refold over the course of one or more unfolding-refolding cycles (Fig. 3B), implying strong interactions with unfolded states. To estimate *K*_D_for PPS binding to partially or fully unfolded states (denoted IU) of PrP, we approximated the system as involving three equilibria: one for ligand binding to N (native state), one for ligand binding to IU, and a third for folding between N and IU. This approximation is very crude because FECs are not equilibrium measurements, it ignores any competition between folding and binding kinetics or force-dependence to the binding/unbinding, and it assumes that the affinity is the same for all partially and fully unfolded states (likely not the case). Under these assumptions, the ratio of FECs starting in IU to those starting in N (whether PPS-bound not), *f*_IU/N_, varies with PPS concentration as

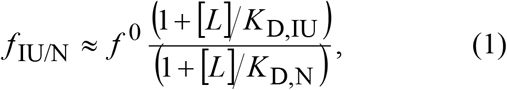

where [*L*] is the ligand concentration, *f*^0^ is the ratio of FECs starting in IU to FECs starting in N in the absence of PPS, *K*_D,IU_is the dissociation constant for PPS binding to IU, and *K*_D,N_is the dissociation constant for PPS binding to N, taken from Ref. 39. Using the N/IU occupancy statistics in Table S2 to solve for *K*_D,IU_at each concentration and taking the geometric mean, we estimated *K*_D,IU_≈ 2 nM. The binding affinity for unfolded states of PrP was therefore ∼100-fold higher than it was for native PrP, indicating surprising avidity for unfolded states that strongly supports the notion suggested previously that interactions with unfolded parts of the protein likely play an important role in anti-prion activity^39,40^.

## DISCUSSION

These results show that by using SMFS to observe ligand-induced changes in the properties of partially and fully unfolded states of a protein, we can characterize in detail how pharmacological chaperones interact with these states and quantify the binding affinity. In the case of PPS binding to PrP, the dissociation for unfolded states was only ∼2 nM, implying affinity about 100-fold higher than for the native state. This difference led to some surprising behavior, such as an increase in the tendency of PrP to show incomplete unfolding (or no unfolding events at all) as the PPS concentration was reduced from 0.5 μM (above *K*_D_for binding the native fold) to 10 nM (well below *K*_D_for binding the native fold but above that for binding unfolded states): tight PPS binding to fully or partially unfolded states increased the likelihood of incomplete (or no) unfolding for [PPS] ≈2 nM, but for binding to N could compete with binding to IU for [PPS] ≈*K*_D,N_, increasing the likelihood of complete folding. Even at the highest concentration tested here, however, the fraction of FECs showing complete unfolding remained significantly below that in the absence of PPS, reflecting the higher affinity of the binding to IU. A similar picture also explains the monotonic increase in unfolding heterogeneity *Δ* with PPS concentration: the increased number of accessible binding sites at higher concentrations raised *Δ*.

Considering what states showed interaction with PPS, we found that PPS was very promiscuous: it not only bound to every state observed in the native folding pathways of BvPrP (*i*.*e*., N, I_1_through I_4_, and U), but also to two additional, mostly-unfolded states (I′_5_and I′_6_). Intriguingly, the latter two states had the same lengths as two of the intermediates seen in the folding pathways for canine PrP^43^, which is naturally immune to propagated misfolding and hence prion disease^50^. However, it is unclear if I′_5_and I′_6_are homologous to the intermediates from CaPrP or some other state, for example involving partial unfolding of the helices in I_3_and I_4_. We speculate that the promiscuity of PPS binding to PrP arises from its high negative charge, which drives electrostatic interactions with positively charged regions of PrP^39,51^, which has a net positive charge at neutral pH. Positive amino acids that are normally buried in the core of the native fold could provide new binding sites when exposed during unfolding, supporting binding to a wide variety of partially or fully unfolded states. The observation that sometimes intermediate states bound to PPS unfolded without difficulty, albeit at higher forces than without PPS (Fig. 5B), whereas at other times they were ‘locked down’ by PPS binding and did not unfold for more than one round of unfolding and refolding (Fig. 5E), suggests that multiple binding modes with differing stability must exist for a given intermediate of PrP, possibly reflecting interactions with different combinations of exposed charges.

The intermediate I_1_at 10 nm was one of the states most frequently observed to interact with PPS (Fig. 5D). Considering the proposed structure of I_1_consisting of all 3 helices but no β-strands^43^, we speculate that PPS binds particularly well to the helix-2/helix-3 interface that is normally covered by strands 1 and 2. Intriguingly, previous work found that deleting the β-strands inhibits conversion of the mutant PrP to PrP^Sc^ and even blocks the conversion of co-expressed wild-type PrP^52^, supporting the notion that inhibition of β-strand formation by PPS binding could be important for the anti-prion activity of PPS.

One particularly notable feature of BvPrP is that it forms long-lived metastable misfolded states in wild-type monomers, unlike other PrP variants, where monomers only form unstable, very short-lived mis-folded states^43–45^. The misfolded conformers in BvPrP form *de novo* from the fully unfolded state, not via partially folded intermediates on the native pathways as in some other proteins^47^. Two of the misfolded conformers observed in BvPrP monomers, M_1_and M_2_, have lengths consistent with the PPS-induced states I′_5_and I′_6_, raising the question of whether I′_5_and I′_6_might be misfolded. Given that M_1_and M_2_form solely and directly from U^43^, any examples where PrP unfolding into I′_5_and I′_6_from a native-like state (N, I_1_to I_3_) as in Fig. 4 can be ruled out as possible incarnations of M_1_and M_2_. Inspection of consecutive pulling cycles, in which the PrP was first completely unfolded, showed that the states I′_5_and I′_6_were not populated following refolding from the completely unfolded state, implying that I′_5_and I′_6_likely do not correspond to M1 or M2. The third mis-folded conformer seen in BvPrP monomers, M_3_, has 8 nm of unfolded contour length, which overlaps with the tail of the I_1_distribution for PPS bound, but again it only forms from the unfolded state. There was no clear evidence for M3 in the FECs with PPS, suggesting that PPS binding inhibits monomer misfolding.

Finally, we note that the approach described here to quantify binding affinity to unfolded states has a number of limitations, including the crude assumptions that the system is in equilibrium and that all partially and fully unfolded states share the same binding affinity. These limitations could, in principle, be overcome by using higher-throughput measurements to analyze the unfolding of each distinct state separately and then applying tools such as the Jarzynski equality or other fluctuation theorems^53,54^ to recover the equilibrium free energy of ligand binding. Such improvements would be technically challenging but provide a more complete and accurate picture of the interactions of ligands like chaperones with unfolded states of proteins.

## METHODS

### Sample Preparation

Recombinant bank vole prion protein (BvPrP) was expressed in *E. coli*, purified, and refolded as described previously^43^. Briefly, the BvPrP construct comprised residues 90−231, with a cysteine residue added to each terminus for attaching DNA handles. Terminal Cys residues were reduced prior to handle conjugation then activated with a 100-fold molar excess of 2,2′-dithiodipyridine (DTDP) at 4°C for 24 h. Excess DTDP was removed by spin filtration, and the activated protein incubated with sulfhydryl-labeled DNA handles at 4°C for 24 h. A 802-bp DNA handle was labeled with biotin and a 2100-bp handle with digoxigenin, enabling specific attachment of the protein-DNA chimera to avidin- and anti-digoxigenin-functionalized polystyrene beads, respectively (diameters 600 nm and 820 nm), to form dumbbell constructs suitable for optical trapping. The resulting dumbbells were diluted to 500 fM in measurement buffer and introduced into the sample chamber for single-molecule force spectroscopy (SMFS). The measurement buffer consisted of 50 mM MOPS (pH 7.0) and 200 mM KCl in the presence of 5kDa PPS at the specified experimental concentrations (0.01 µM, 0.1 µM and 0.5 µM); it also contained an oxygen-scavenging system comprising 8 mU/µL glucose oxidase, 20 mU/µL catalase, and 0.01% w/v D-glucose. No accessory proteins were included in the molecular construct used for SMFS. Fresh protein–DNA constructs were prepared regularly to minimize aggregation and reduce the occurrence of multiple tethers during SMFS measurements.

### SMFS measurements

SMFS measurements were performed using a custom-built dual-beam optical tweezers instrument following previously established procedures^43^. Briefly, individual PrP dumbbells were captured between two independently controlled optical traps with stiffnesses of 0.43 and 0.56 pN/nm. The traps were moved apart at a constant speed of 200 nm/s to generate FECs. Between successive unfolding–refolding cycles, the molecule was held at near-zero force for 5 s to allow refolding before the next pulling cycle. Force and extension data were acquired at 20 kHz, filtered online using an 8-pole Bessel filter at the Nyquist frequency. To minimize the effects of instrumental drift during extended measurements, position sensors were recalibrated after every 100 pulls. Only traces consistent with single-molecule tethering (via the persistence length measured in FECs) were included in the analysis. Measurements in presence of PPS from 14 BvPrP molecules yielded a total of ∼3,750 FECs. Measurements without PPS were reported previously^43^.

### Contour length analysis

To quantify the changes in contour length (Δ*L*_*c*_) during unfolding of the protein, the FECs were fitted to an extensible WLC model as described previously^43^. The force-extension relationship was modelled as:

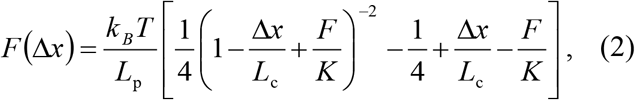

where *L*_p_is the polymer persistence length, *L*_c_is the contour length, and *K* the elastic modulus. The protein-DNA construct was represented by two WLC elements in series, corresponding to the DNA handles and the unfolded protein, respectively. The parameters *L*_c_, *L*_p_, and *K* of the DNA handles were treated as free parameters and determined by fitting the fully folded portion of each FEC and were subsequently fixed for fitting intermediate and unfolded regions.

For the protein component, *L*_p_and *K* were fixed at 0.65 nm and 2,000 pN, respectively, while *L*_c_was allowed to vary and was used as the sole fitting parameter for each structural state. The contour length changes obtained from individual FEC branches were used to construct cumulative Δ*L*_c_distributions relative to the fully folded state. For each BvPrP molecule, the resulting distributions were comparable, indicating consistent folding behavior across molecules; measurements from all molecules were combined for subsequent analysis.

### Unfolding force analysis

The unfolding force distributions, *p*(*F*), were fit to a model that involves heterogeneous energy barriers with multiple initial states that can interconvert^49^, given by

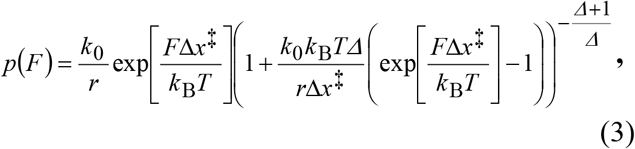

where *k*_0_is the zero-force unfolding rate averaged over the ensemble of initial states, Δ*x*^‡^ is the average distance to the transition state from the ensemble of starting states, and *Δ* is a dimensionless parameter reflecting the extent of heterogeneity (with larger *Δ* indicating greater heterogeneity).

## Acknowledgements

This work was funded by the Alberta Prion Research Institute (grant numbers 201600014, 201800008) and Canadian Institutes of Health Research (PJT-185931).

## Author contributions

SP and MTW designed research; CRG provided reagents; SP made measurements; SP analyzed data; SP and MTW wrote manuscript.

## Supplementary information

**Fig. S1.**
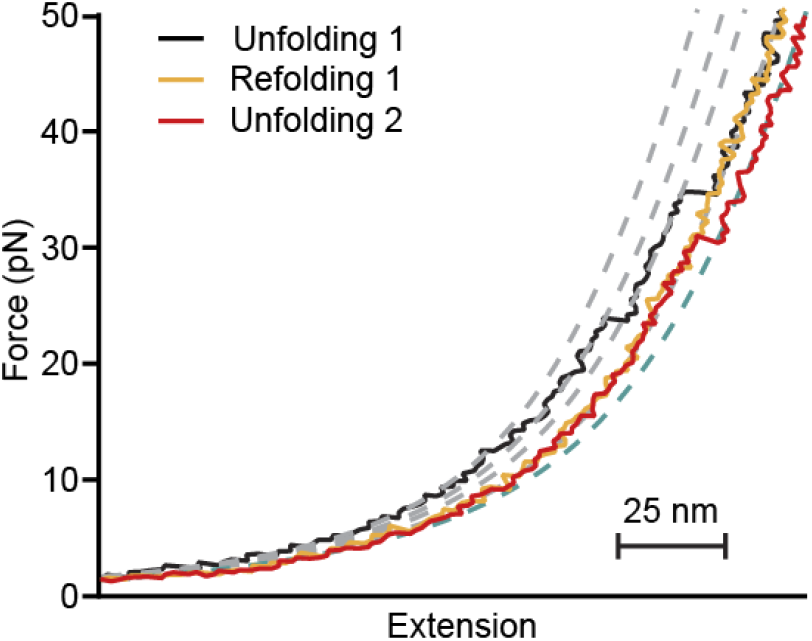
Sequential FECs showing an unfolding-refolding-unfolding cycle where BvPrP did not refold but started the second unfolding curve in the same intermediate where it ended the previous unfolding curve.

**Table S1.** The mean Δ*L*_c_ observed for intermediate states with respect to the native state and the corresponding mean unfolding force, in presence of PPS.

| Intermediate state | Mean unfolding $\Delta L_c$ (nm) | Mean unfolding force (pN) |
| --- | --- | --- |
| $I_1$ | $9.8 \pm 0.2$ | $15.4 \pm 0.3$ |
| $I_2$ | $14.1 \pm 0.2$ | $20.5 \pm 0.7$ |
| $I_3$ | $18.4 \pm 0.2$ | $22.9 \pm 0.7$ |
| $I_4$ | $24.4 \pm 0.2$ | $27 \pm 1$ |
| $I'_5$ | $22.1 \pm 0.2$ | $26.2 \pm 0.8$ |
| $I'_6$ | $27.9 \pm 0.2$ | $29.4 \pm 0.9$ |

**Table S2.** The fraction of occupancy of N and I/U for different concentrations of PPS.

| PPS concentration ($\mu\text{M}$) | Fraction in N | Fraction in I/U |
| --- | --- | --- |
| 0.01 | 0.21 | 0.79 |
| 0.1 | 0.31 | 0.69 |
| 0.5 | 0.36 | 0.64 |

## References

1. Louros N, Schymkowitz J, Rousseau F. Mechanisms and pathology of protein misfolding and aggregation. Nat. Rev. Mol. Cell. Biol. 24: 912–33 (2023).

2. Devi S, Chaturvedi M, Fatima S, Priya S. Environmental factors modulating protein conformations and their role in protein aggregation diseases. Toxicology 465: 153049 (2022).

3. Speer SL, Stewart CJ, Sapir L, Harries D, Pielak GJ. Macromolecular crowding is more than hard-core repulsions. Annu. Rev. Biophys. 51: 267–300 (2022).

4. Tokuriki N, Tawfik DS. Stability effects of mutations and protein evolvability. Curr. Opin. Struct. Biol. 19: 596–604 (2009).

5. Hartl FU, Bracher A, Hayer-Hartl M. Molecular chaperones in protein folding and proteostasis. Nature 475: 324–332 (2011).

6. Stebbins CE, Russo AA, Schneider C, Rosen N, Hartl FU, Pavletich NP. Crystal structure of an Hsp90-geldanamycin complex: targeting of a protein chaperone by an antitumor agent. Cell 89: 239–250 (1997).

7. Clerico EM, Tilitsky JM, Meng W, Gierasch LM. How hsp70 molecular machines interact with their substrates to mediate diverse physiological functions. J. Mol. Biol. 427: 1575–1588 (2015).

8. Mayer MP, Gierasch LM. Recent advances in the structural and mechanistic aspects of Hsp70 molecular chaperones. J. Biol. Chem. 294: 2085–2097 (2019).

9. Fan JQ, Ishii S, Asano N, Suzuki Y. Accelerated transport and maturation of lysosomal alpha-galactosidase A in Fabry lymphoblasts by an enzyme inhibitor. Nat. Med. 5: 112–115 (1999).

10. Parenti G, Moracci M, Fecarotta S, Andria G. Pharmacological chaperone therapy for lysosomal storage diseases. Future Med. Chem. 6: 1031–1045 (2014).

11. Chiti F, Kelly JW. Small molecule protein binding to correct cellular folding or stabilize the native state against misfolding and aggregation. Curr. Opin. Struct. Biol. 72: 267–278 (2022).

12. Hammoudeh DI, Follis AV, Prochownik EV, Metallo SJ. Multiple independent binding sites for small-molecule inhibitors on the oncoprotein c-Myc. J. Am. Chem. Soc. 131: 7390–7401 (2009).

13. Ehrnhoefer DE, Bieschke J, Boeddrich A, Herbst M, Masino L, Lurz R, et al. EGCG redirects amyloidogenic polypeptides into unstructured, off-pathway oligomers. Nat. Struct. Mol. Biol. 15: 558–566 (2008).

14. Boeckler FM, Joerger AC, Jaggi G, Rutherford TJ, Veprintsev DB, Fersht AR. Targeted rescue of a destabilized mutant of p53 by an in silico screened drug. Proc. Natl. Acad. Sci. USA 105: 10360–10365 (2008).

15. Bulawa CE, Connelly S, DeVit M, Wang L, Weigel C, Fleming JA, et al. Tafamidis, a potent and selective transthyretin kinetic stabilizer that inhibits the amyloid cascade. Proc. Natl. Acad. Sci. USA 109: 9629–9634 (2012).

16. Dobrev VS, Fred LM, Gerhart KP, Metallo SJ. Characterization of the binding of small molecules to intrinsically disordered proteins. Methods Enzymol. 611: 677–702 (2018).

17. Heller GT, Aprile FA, Vendruscolo M. Methods of probing the interactions between small molecules and disordered proteins. Cell. Mol. Life Sci. 74: 3225–3243 (2017).

18. Chowdhury A, Nettels D, Schuler B. Interaction dynamics of intrinsically disordered proteins from single-molecule spectroscopy. Annu. Rev. Biophys. 52: 433–462 (2023).

19. Petrosyan R, Narayan A, Woodside MT. Single-molecule force spectroscopy of protein folding. J. Mol. Biol. 433:167207 (2021).

20. Woodside MT, Block SM. Reconstructing folding energy landscapes by single-molecule force spectroscopy. Annu. Rev. Biophys. 43: 19–39 (2014).

21. Mashaghi A, Kramer G, Bechtluft P, Zachmann-Brand B, Driessen AJM, Bukau B, et al. Reshaping of the conformational search of a protein by the chaperone trigger factor. Nature. 500: 98–101 (2013).

22. Perales-Calvo J, Giganti D, Stirnemann G, Garcia-Manyes S. The force-dependent mechanism of DnaK-mediated mechanical folding. Sci. Adv. 4: eaaq0243 (2018).

23. Ainavarapu SRK, Li L, Badilla CL, Fernandez JM. Ligand binding modulates the mechanical stability of dihydrofolate reductase. Biophys. J. 89: 3337–3344 (2005).

24. Stigler J, Rief M. Calcium-dependent folding of single calmodulin molecules. Proc. Natl. Acad. Sci. USA. 109: 17814–17819 (2012).

25. Colby DW, Prusiner SB. Prions. Cold Spring Harb. Perspect. Biol. 3: a006833 (2011).

26. Caughey B, Baron GS, Chesebro B, Jeffrey M. Getting a grip on prions: Oligomers, amyloids, and pathological membrane interactions. Annu. Rev. Biochem. 78: 177–204 (2009).

27. Vaquer-Alicea J, Diamond MI. Propagation of protein aggregation in neurodegenerative diseases. Annu. Rev. Biochem. 88: 785–810 (2019).

28. Sim VL. Prion disease: Chemotherapeutic strategies. Infect. Disord. - Drug Targets 12: 144–160 (2012).

29. Teruya K, Doh-Ura K. Insights from therapeutic studies for PrP prion disease. Cold Spring Harb. Perspect. Med. 7: a024430 (2017).

30. Sweeney P, Park H, Baumann M, Dunlop J, Frydman J, Kopito R, et al. Protein misfolding in neurodegenerative diseases: implications and strategies. Transl. Neurodegener. 6: 6 (2017).

31. Giles K, Olson SH, Prusiner SB. Developing therapeutics for PrP prion diseases. Cold Spring Harb. Perspect. Med. 7: a023747 (2017).

32. Frei JA, Reidenbach AG, Xu LMH, Gopalakrishnan RM, Casalana D, Sprague DA, et al. Phenotypic screening for small molecules that lower PrP in cultured cells. Mol. Cell Neurosci. 138: 104111 (2026).

33. Sabzehei S, Innocenti N, Rigoli M, Requena JR, Biasini E. Unfolding prion misfolding and the challenge of identifying effective therapeutics: is there hope on the horizon? Expert Opin. Drug Discov. 21: 793–801 (2026).

34. Kamatari YO, Hayano Y, Yamaguchi KI, Hosokawa-Muto J, Kuwata K. Characterizing antiprion compounds based on their binding properties to prion proteins: Implications as medical chaperones. Prot. Sci. 22: 22–34 (2013).

35. Barreca ML, Iraci N, Biggi S, Cecchetti V, Biasini E. Pharmacological agents targeting the cellular prion protein. Pathogens 7: 27 (2018).

36. Staderini M, Vanni S, Baldeschi AC, Giachin G, Zattoni M, Celauro L, et al. Bifunctional carbazole derivatives for simultaneous therapy and fluorescence imaging in prion disease murine cell models. Eur. J. Med. Chem. 245: 114923 (2023).

37. Stincardini C, Massignan T, Biggi S, Elezgarai SR, Sangiovanni V, Vanni I, et al. An antipsychotic drug exerts anti-prion effects by altering the localization of the cellular prion protein. PLoS One 12: e0182589 (2017).

38. Vogtherr M, Grimme S, Elshorst B, Jacobs DM, Fiebig K, Griesinger C, et al. Antimalarial drug quinacrine binds to C-terminal helix of cellular prion protein. J. Med. Chem. 46: 3563–3564 (2003).

39. Petrosyan R, Patra S, Rezajooei N, Garen CR, Woodside MT. Unfolded and intermediate states of PrP play a key role in the mechanism of action of an antiprion chaperone. Proc. Natl. Acad. Sci. USA 118: e2010213118 (2021).

40. Gupta AN, Neupane K, Rezajooei N, Cortez LM, Sim VL, Woodside MT. Pharmacological chaperone reshapes the energy landscape for folding and aggregation of the prion protein. Nat. Commun. 7: 12058 (2016).

41. Gupta S, Rao AR, Varadwaj PK, D. S, Mohapatra T. Extrapolation of inter domain communications and substrate binding cavity of camel HSP70 1A: A molecular modeling and dynamics simulation study. PLoS One 10: e0136630 (2015).

42. Watts JC, Giles K, Patel S, Oehler A, DeArmond SJ, Prusiner SB. Evidence that bank vole PrP is a universal acceptor for prions. PLoS Pathog. 10: e1003990 (2014).

43. Anand U, Patra S, Sekar RV, Garen CR, Woodside MT. Different folding mechanisms in prion proteins from mammals with different disease susceptibility observed at the singlemolecule level. Proc. Natl. Acad. Sci. USA 122: e2416191122 (2025).

44. Yu H, Liu X, Neupane K, Gupta AN, Brigley AM, Solanki A, et al. Direct observation of multiple misfolding pathways in a single prion protein molecule. Proc. Natl. Acad. Sci. USA 109: 5283–5288 (2012).

45. Yu H, Dee DR, Liu X, Brigley AM, Sosova I, Woodside MT. Protein misfolding occurs by slow diffusion across multiple barriers in a rough energy landscape. Proc. Natl. Acad. Sci. USA 112: 8308–8313 (2015).

46. Christen B, Pérez DR, Hornemann S, Wüthrich K. NMR structure of the bank vole prion protein at 20 °C contains a structured loop of residues 165–171. J. Mol. Biol. 383: 306–312 (2008).

47. Sen Mojumdar S, Scholl ZN, Dee DR, Rouleau L, Anand U, Garen C, et al. Partially native intermediates mediate misfolding of SOD1 in single-molecule folding trajectories. Nat. Commun. 8: 1881 (2017).

48. Zhang Y, Dudko OK. A transformation for the mechanical fingerprints of complex biomolecular interactions. Proc. Natl. Acad. Sci. USA 110: 16432–16437 (2013).

49. Hinczewski M, Hyeon C, Thirumalai D. Directly measuring single-molecule heterogeneity using force spectroscopy. Proc. Natl. Acad. Sci. USA 113: E3852–E3861 (2016).

50. Fernández-Borges N, Parra B, Vidal E, Eraña H, Sánchez-Martín MA, de Castro J, et al. Unraveling the key to the resistance of canids to prion diseases. PLoS Pathog. 13: e1006716 (2017).

51. Taubner LM, Bienkiewicz EA, Copié V, Caughey B. Structure of the flexible amino-terminal domain of prion protein bound to a sulfated glycan. J. Mol. Biol. 395: 475–490 (2010).

52. Vorberg I, Chan K, Priola SA. Deletion of β-strand and α-helix secondary structure in normal prion protein inhibits formation of its protease-resistant isoform. J. Virol. 75: 10024–10032 (2001).

53. Jarzynski C. Nonequilibrium equality for free energy differences. Phys. Rev. Lett. 78: 2690 (1997).

54. Camunas-Soler J, Alemany A, Ritort F. Experimental measurement of binding energy, selectivity, and allostery using fluctuation theorems. Science 355: 412–415 (2017).

